## Supplementary figures and images for "Mitochondrial biogenesis is transcriptionally repressed in lysosomal lipid storage diseases"

### Figure 1 - supplement 1

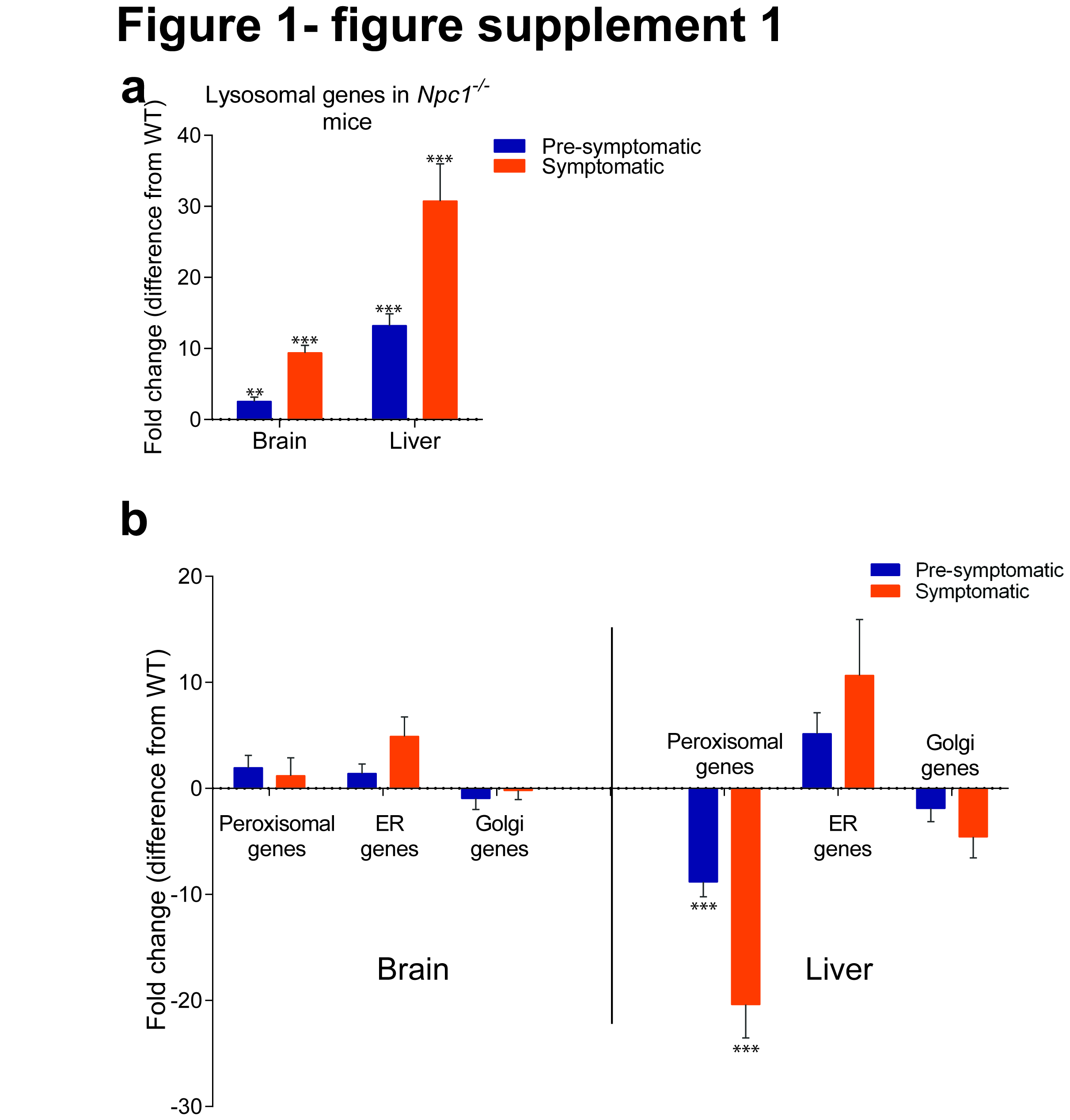

### Figure 1 - supplement 2

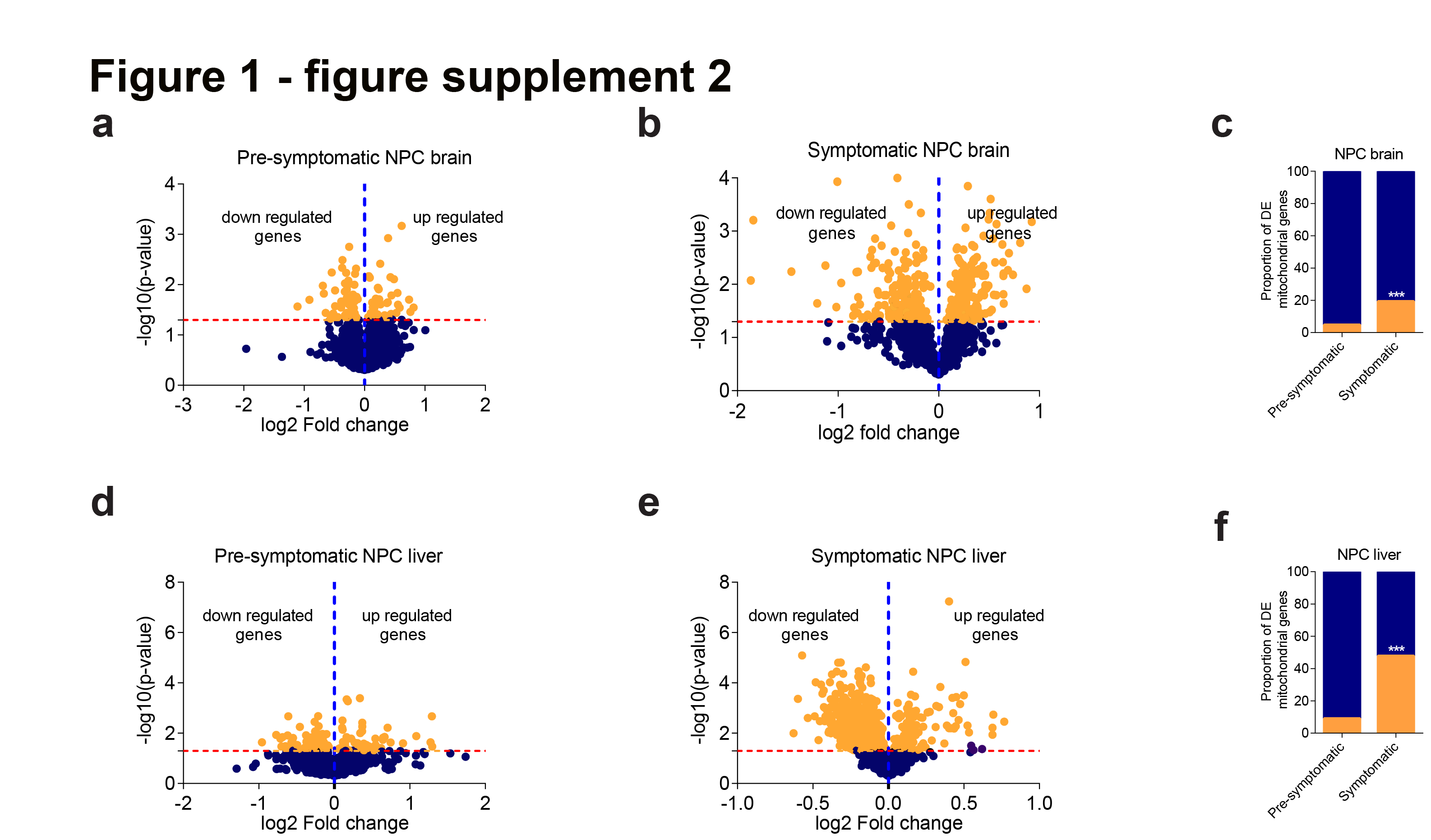

### Figure 3 - supplement 1

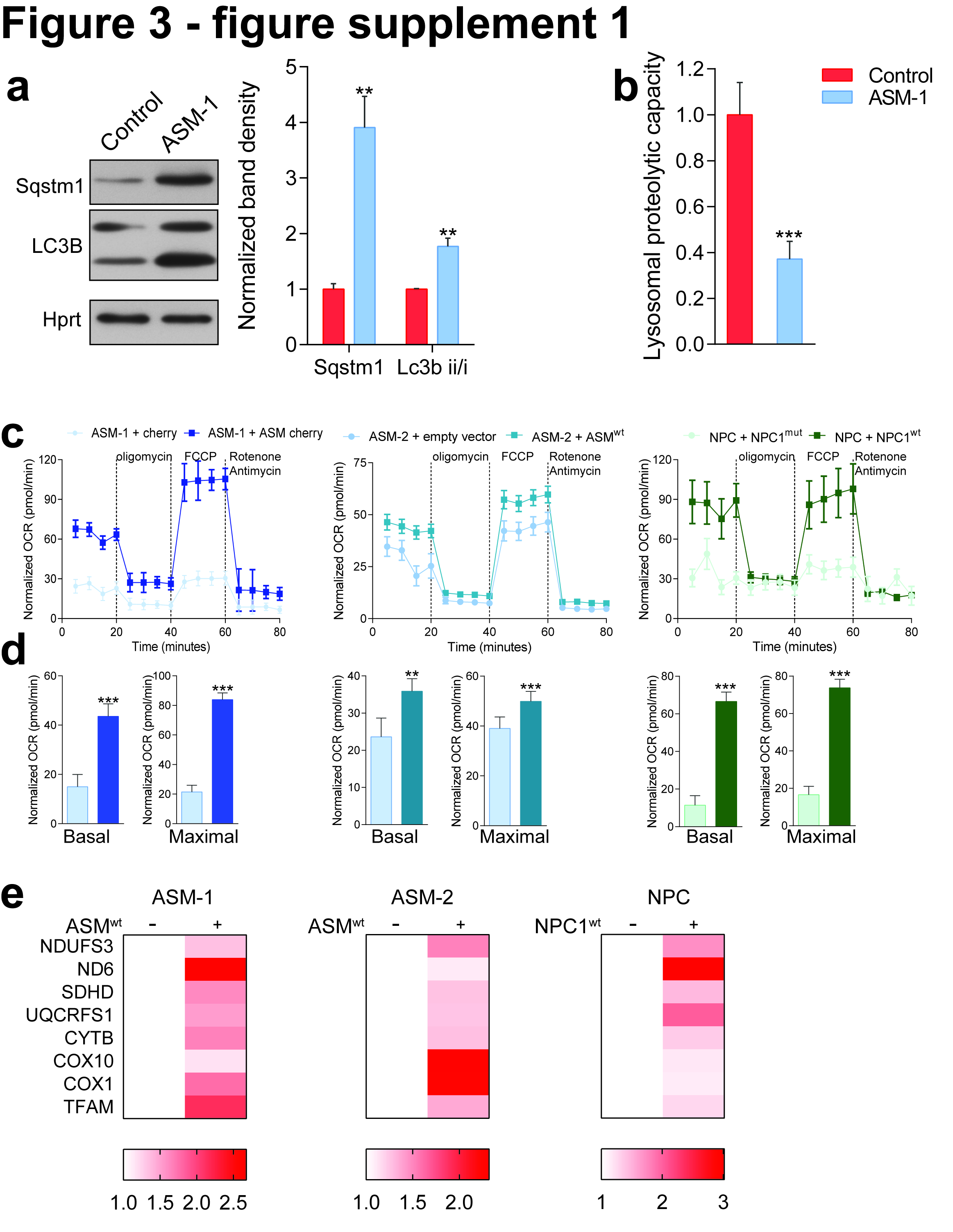

### Figure 3 - supplement 2

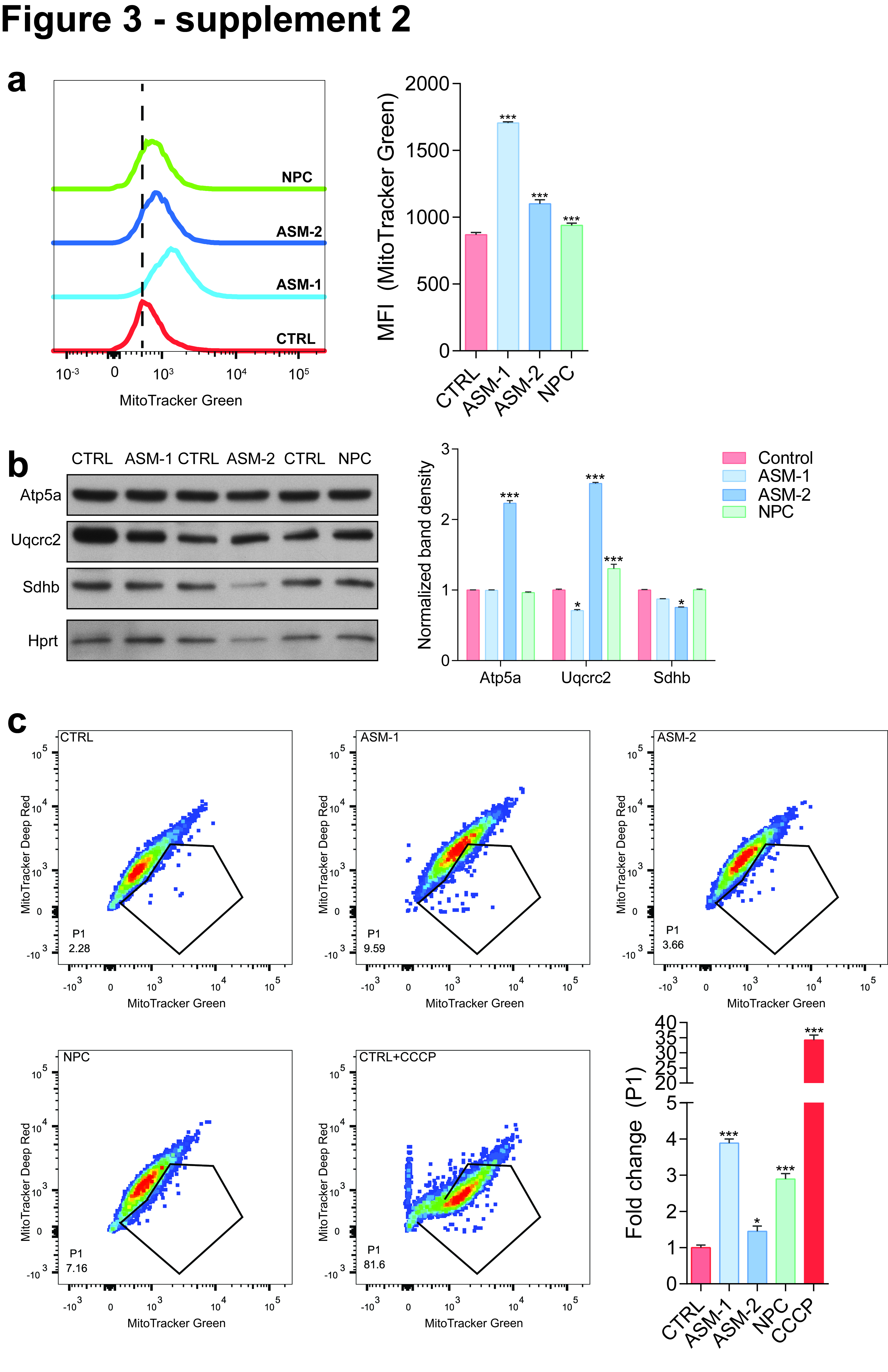

### Figure 3 - supplement 3

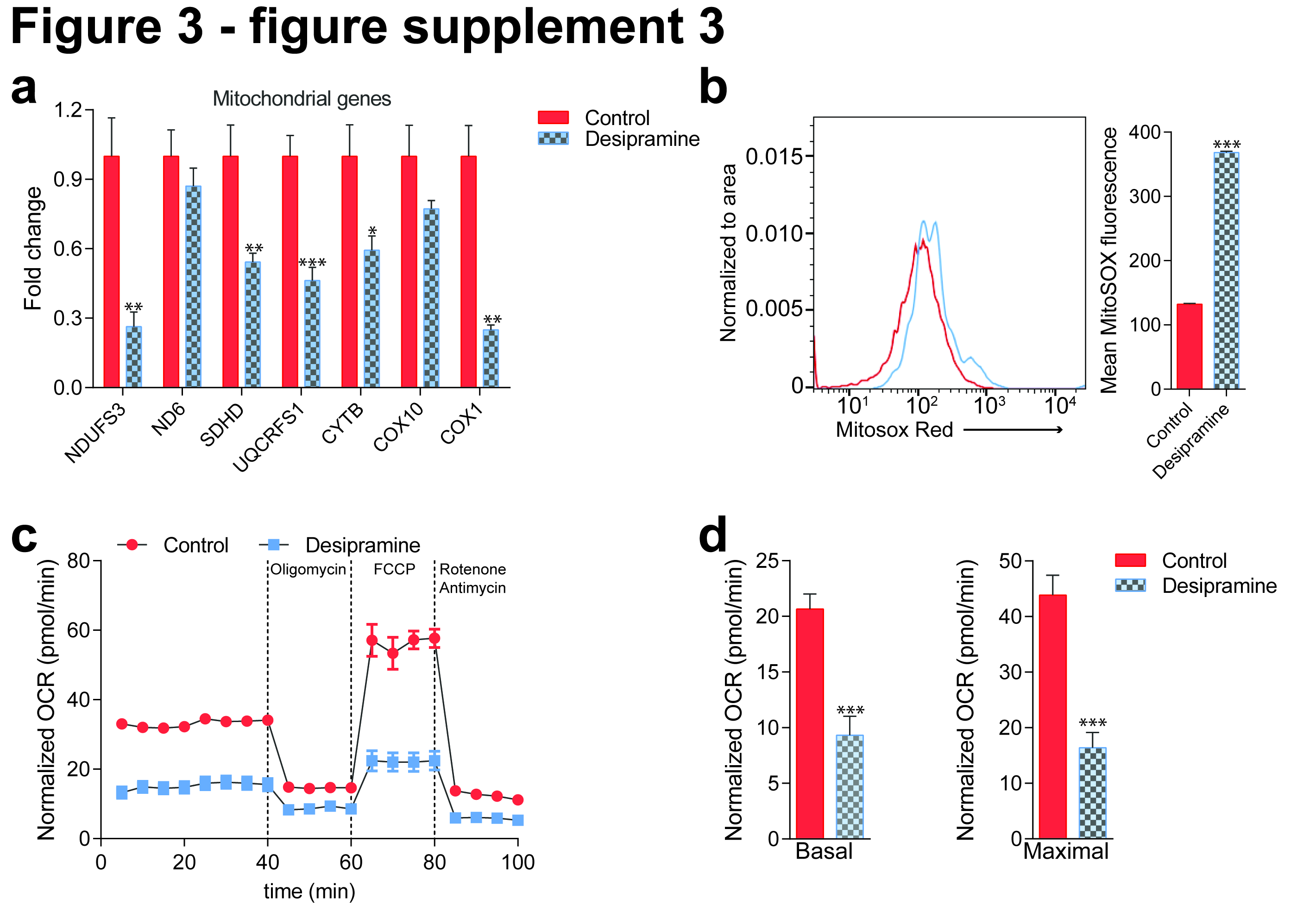

### Figure 3 - supplement 4

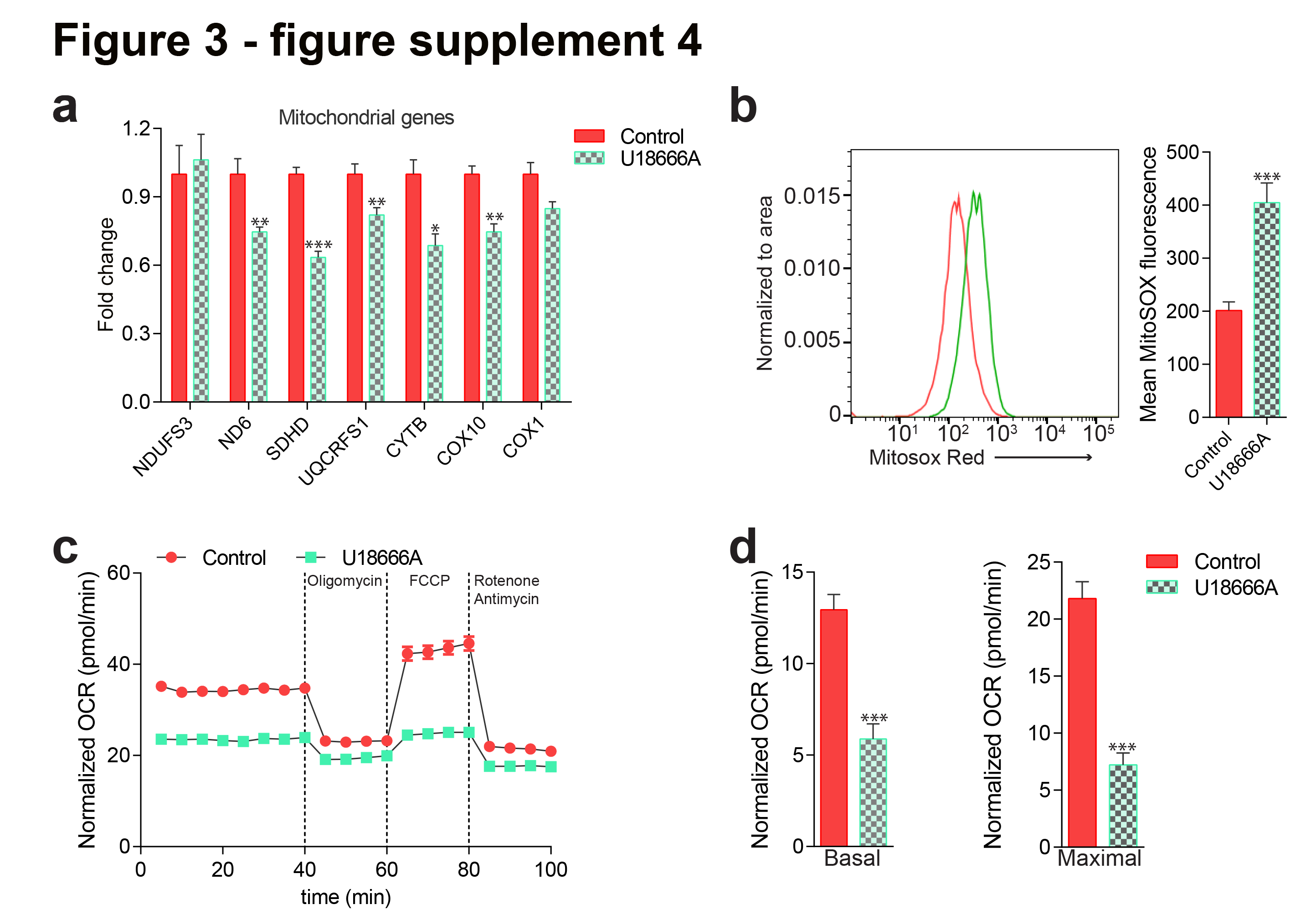

### Figure 4 - supplement 1

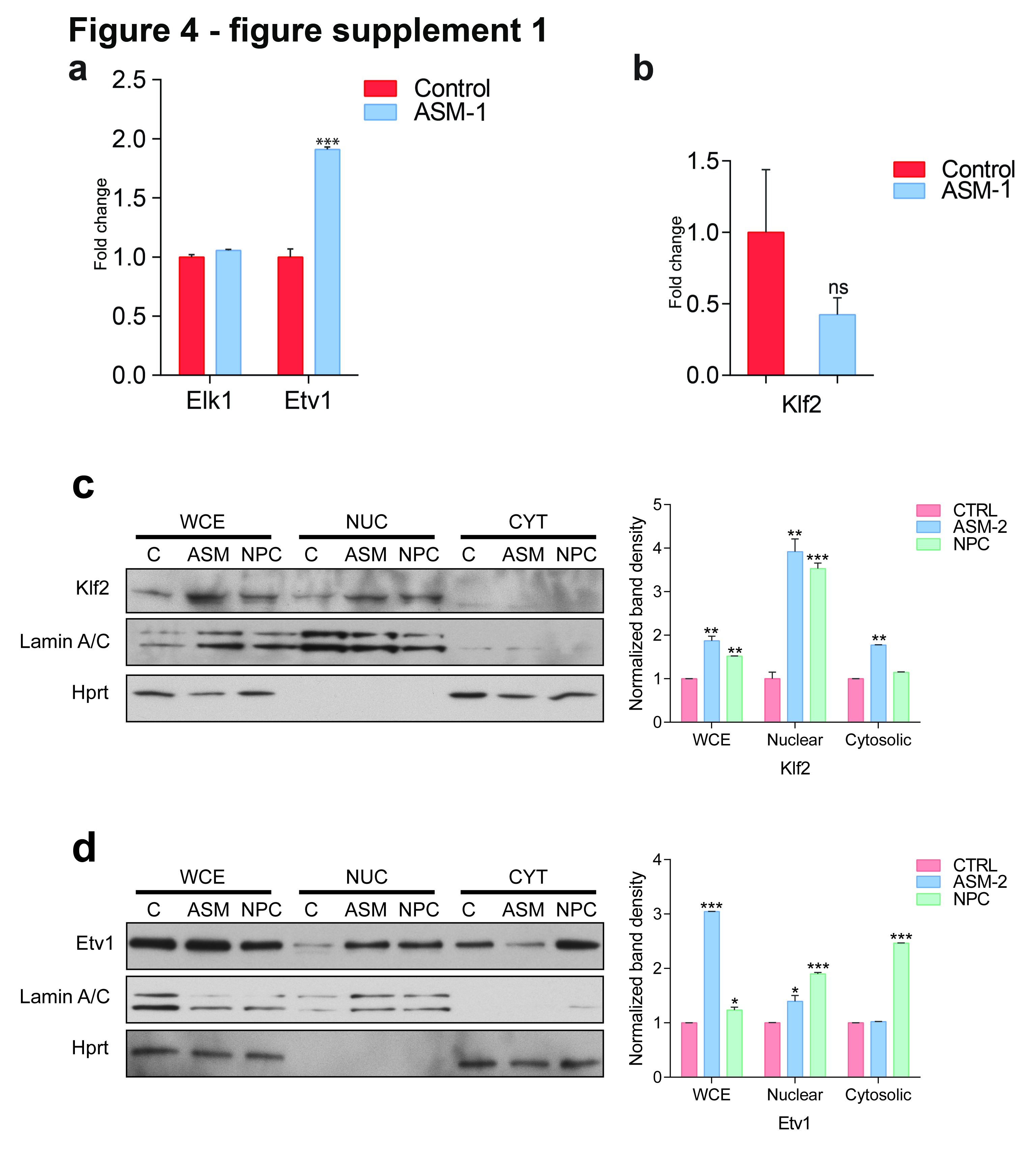

### Figure 4 - supplement 2

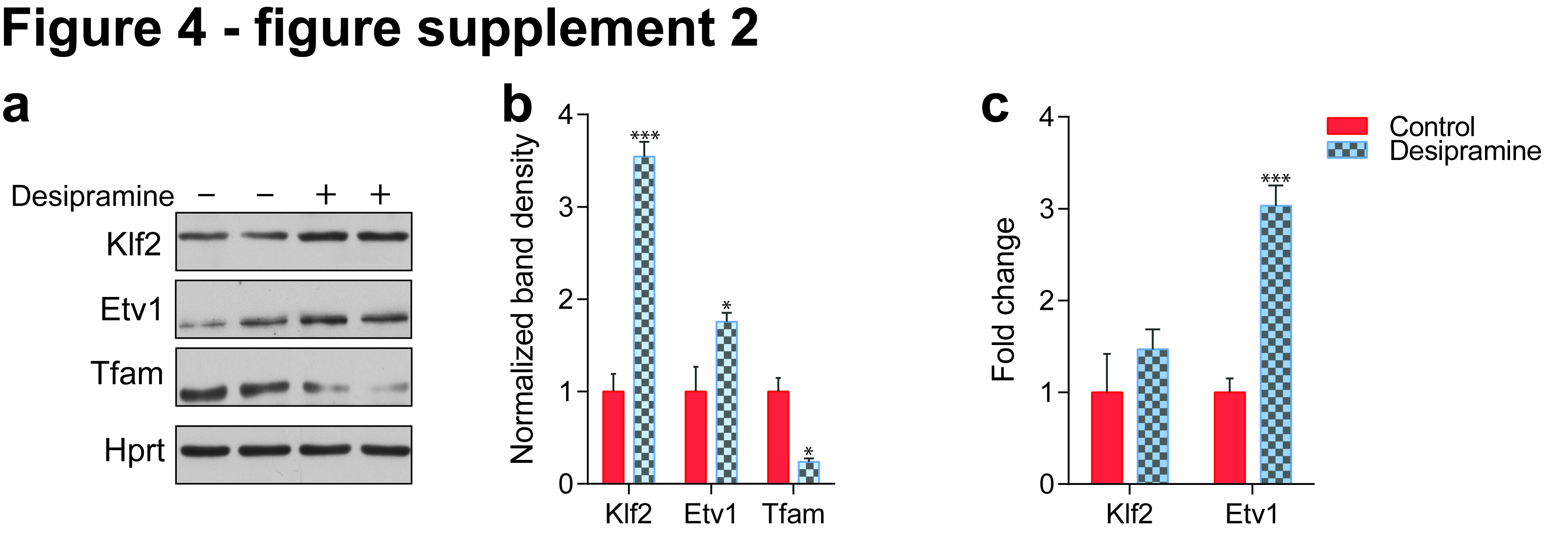

### Figure 4 - supplement 3

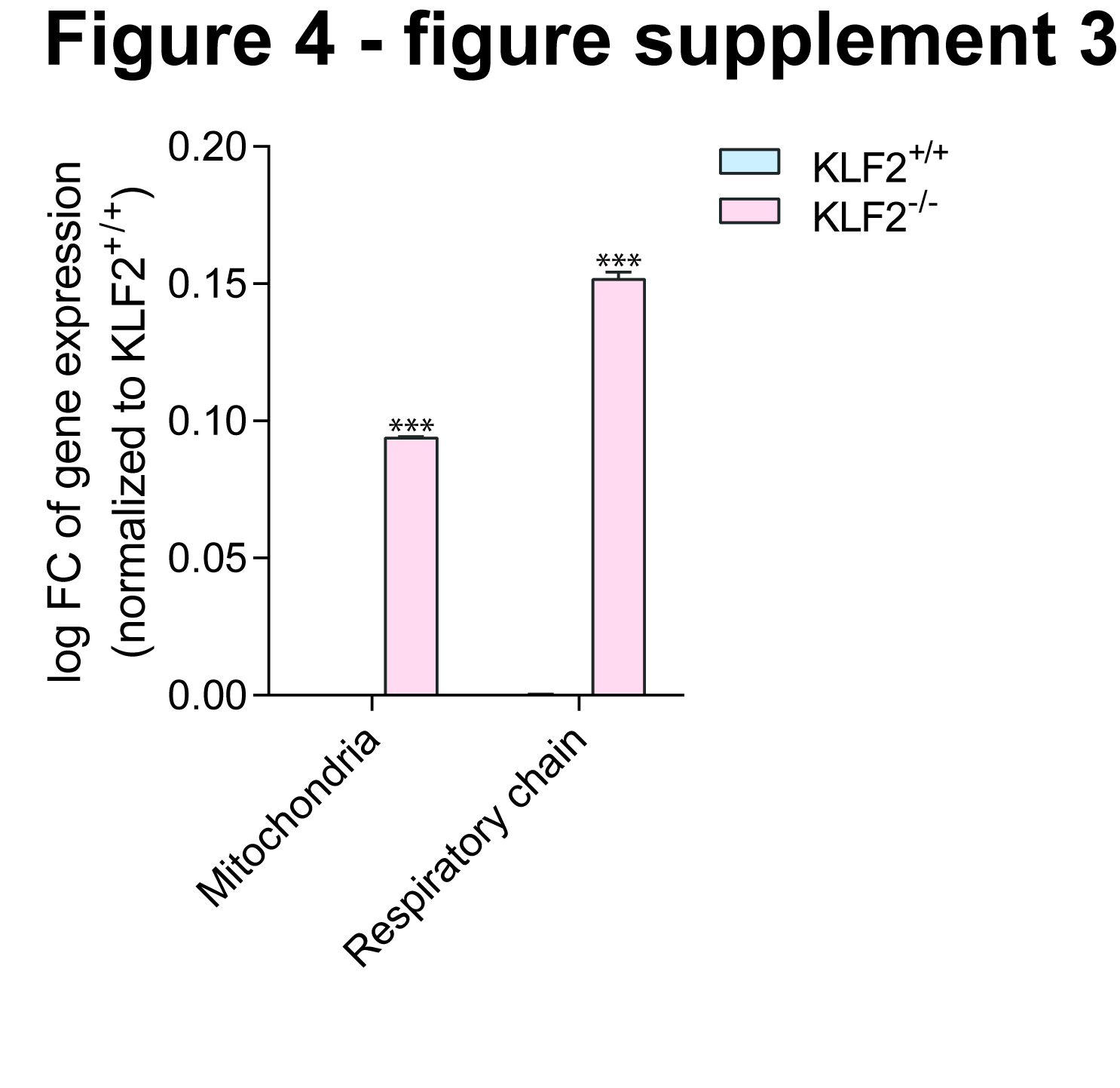

### Figure 4 - supplement 4

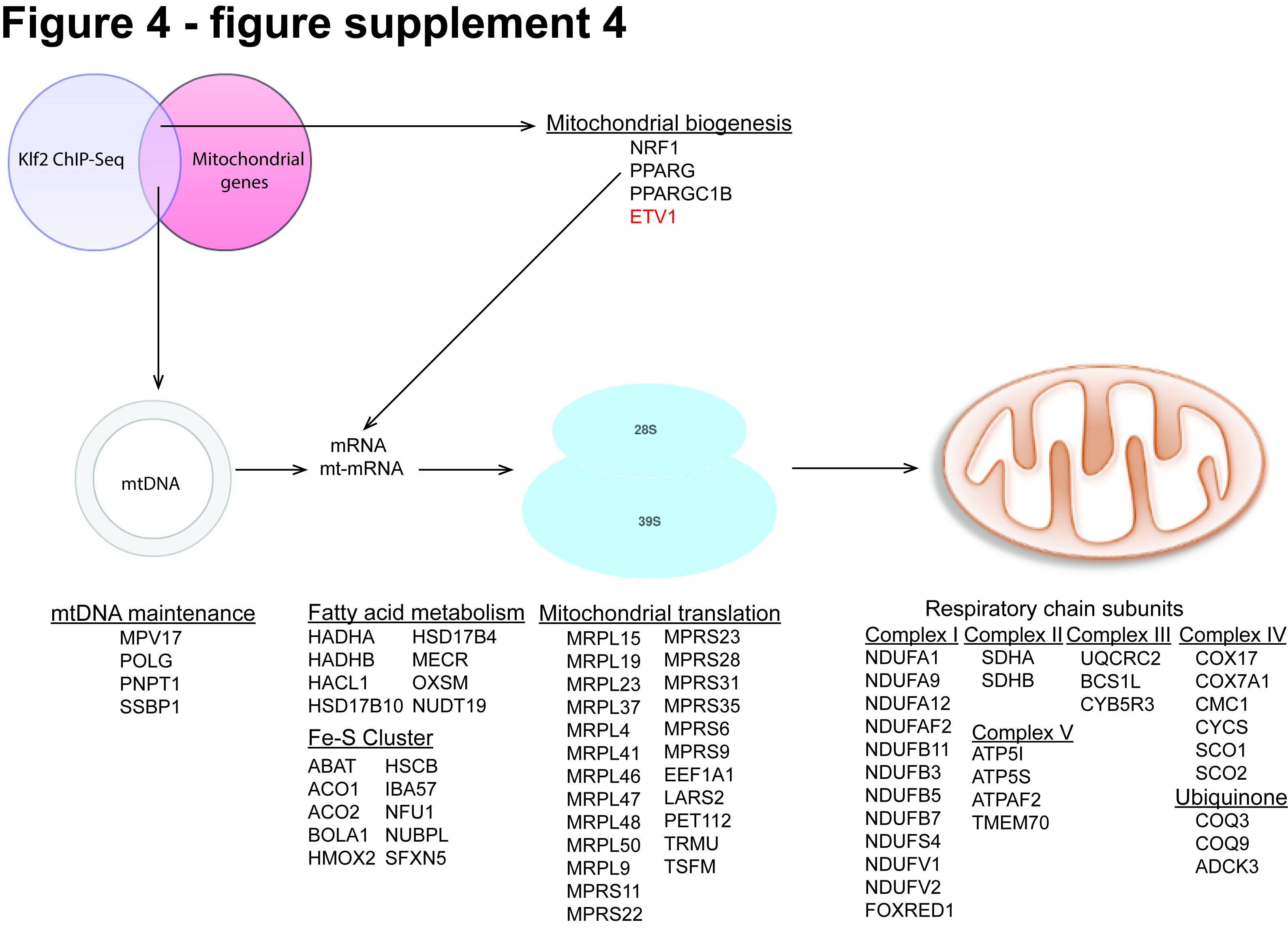

### Figure 4 - supplement 5

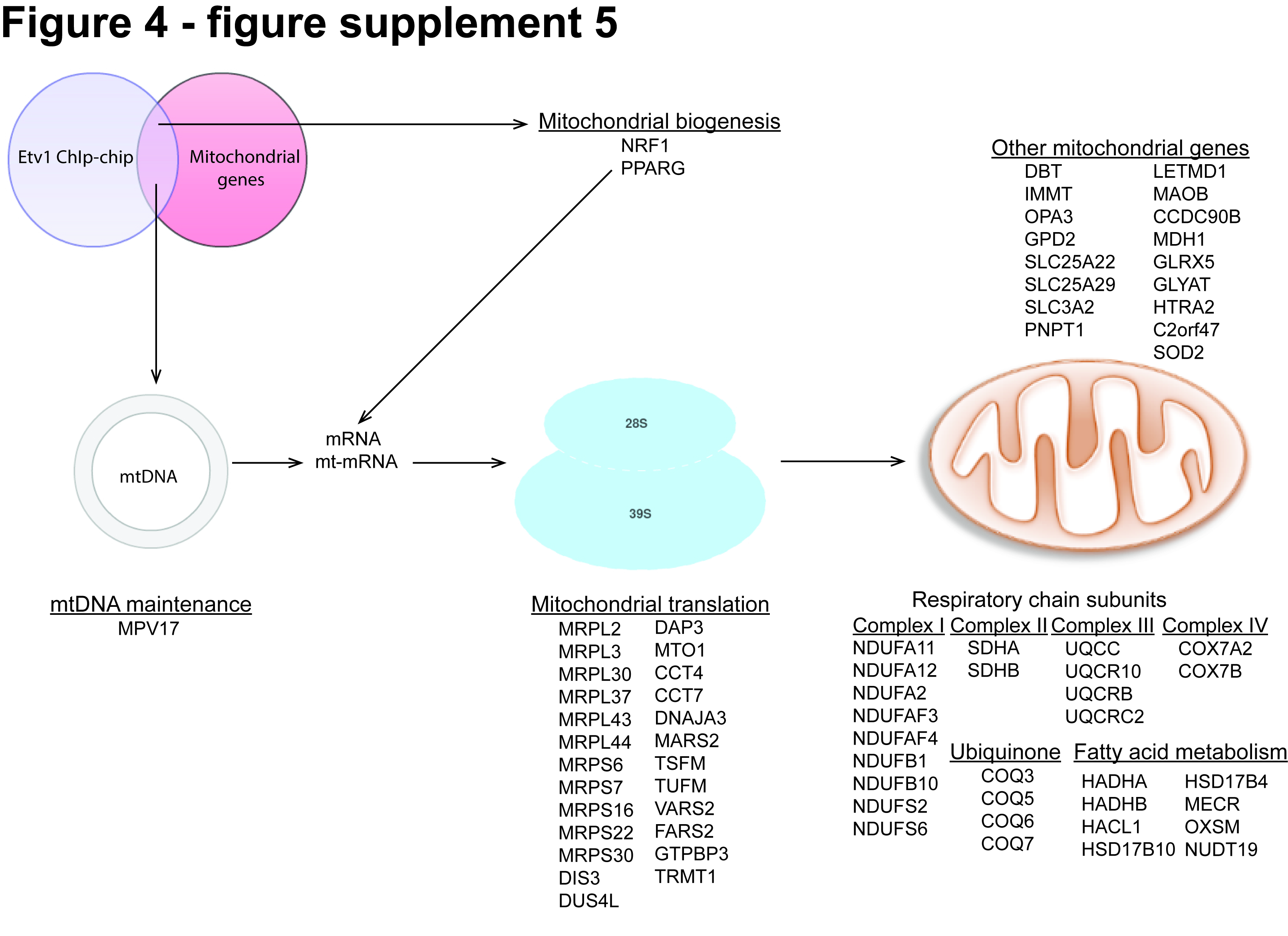

### Figure 5 - supplement 1

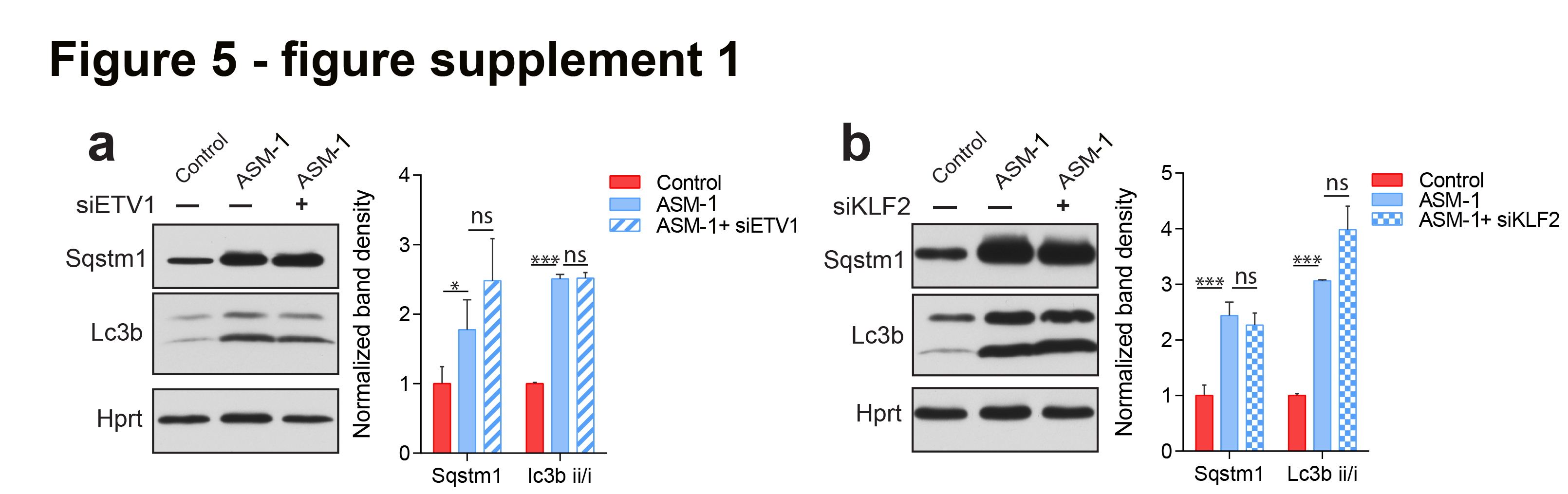

### Figure 7 - supplement 1

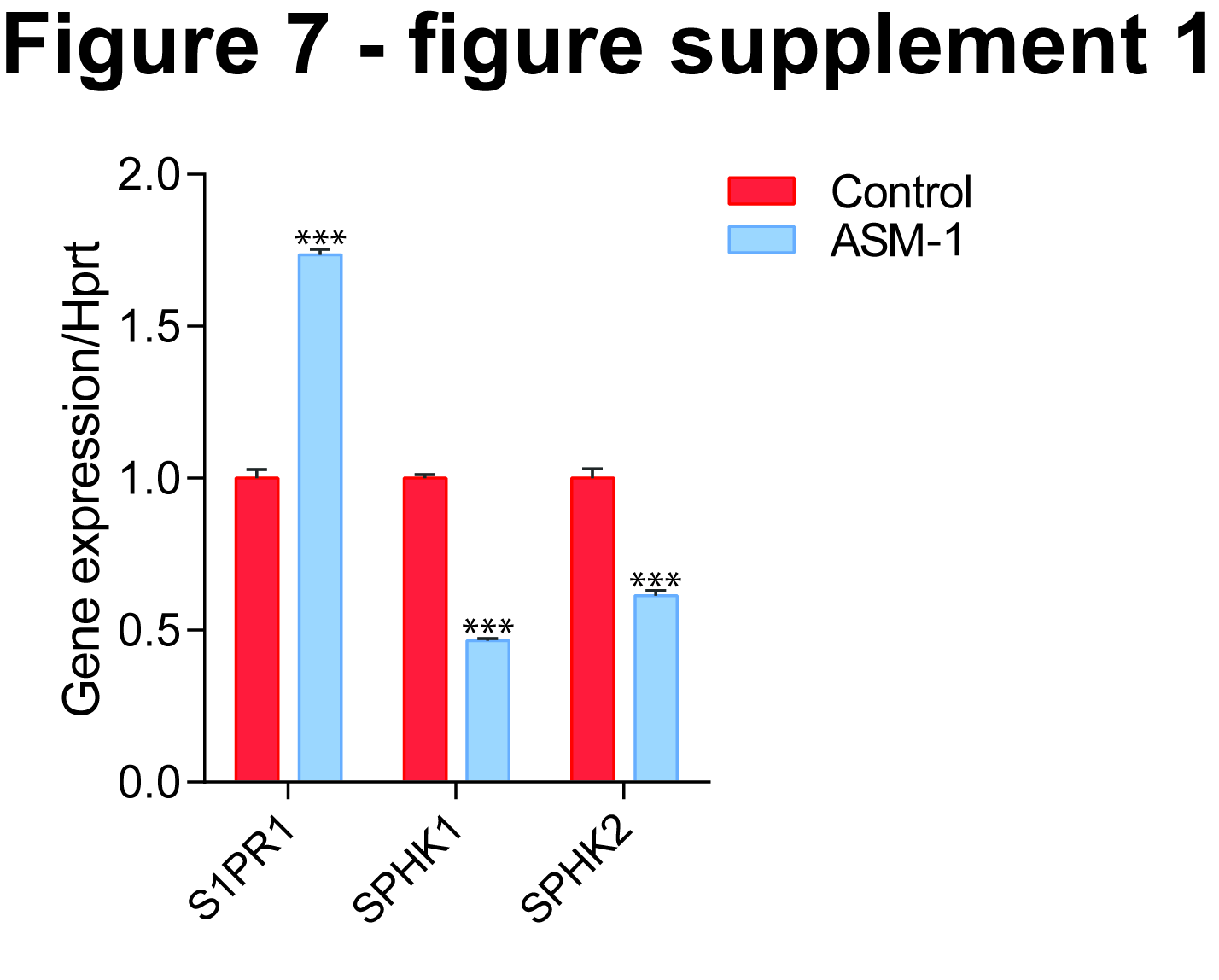
